# Wormtrails: a python package for viewing *C. elegans* movement

**DOI:** 10.1101/2025.11.14.688522

**Authors:** Christopher Dante Ashih, Yuyan Xu, Javier Apfeld

**Affiliations:** Biology Department, Northeastern University, Boston, Massachusetts, United States

## Abstract

Wormtrails is a python package designed to create images and videos depicting the motion of *C. elegans* on solid media. Darkfield or brightfield images may be converted into stills or movies, with time encoded by color. This package was primarily designed to be used for visualizing locomotion during chemotaxis, but it may be used to visualize locomotory patterns in a wide array of behavioral assays.

## Results and discussion

Wide field microscopy recordings of *C. elegans* on solid media are commonly used to document and share experimental results (Breimann et al., 2019). Although many high-throughput tracking and analysis tools are available (Baek et al., 2002; Churgin et al., 2017; Javer et al., 2018; Ramot et al., 2008; Roussel et al., 2007; Stroustrup et al., 2013; Swierczek et al., 2011), behavior is most often presented either as raw images or video recordings (Cowen et al., 2024; Tasnim et al., 2025) or as plots of measurables such as position and speed (Matsumura et al., 2025; Schiffer et al., 2021). Yet, despite their value, these formats are often not well suited to the initial exploration of a dataset. Exploration is critical because it can reveal unexpected patterns and heterogeneity in behavior which can guide the subsequent choice of meaningful summary metrics and hypotheses to test. Inspired by Harold Edgerton’s multiflash photography (Edgerton, 1959) and long exposure imaging, the wormtrails Python package provides fast, user-friendly post-processing to visualize recent locomotory patterns. Wormtrails complements existing quantitative pipelines by facilitating the early, exploratory inspection of behavioral data.

In the simplest case, wormtrails can display the full set of tracks accumulated over an entire recording (Figure 1A). This produces visualizations analogous to oblique illumination of tracks left on solid media (Mori & Ohshima, 1995) or to plotting all recorded positions acquired by tracking (Luo et al., 2014), depending on whether a dark or bright background is used. In a single still image, complex behaviors such as dwelling and roaming can become apparent. For long recordings or dense populations (Figures 1C,D), extensive track overlap can make these visualizations difficult to interpret. To emphasize recent behavior rather than the full positional history, wormtrails can construct “track frames” from a shorter temporal window (Figures 1B,D). Applying a fixed frame window across the recording yields a video in which each frame shows only the recent paths taken (Videos 1-6). This sliding-window approach helps declutter recordings with many worms or long assays and provides a clearer representation of each worm’s recent locomotory behavior. Temporal information can be further encoded using color to indicate when an animal occupied a given position. With short time-frame windows, a simple fade-in/fade-out scheme allows older portions of the track to gradually disappear rather than stopping abruptly, revealing the recent direction of movement (Figures 1B,D, Videos 3,6).

**Figure 1.**
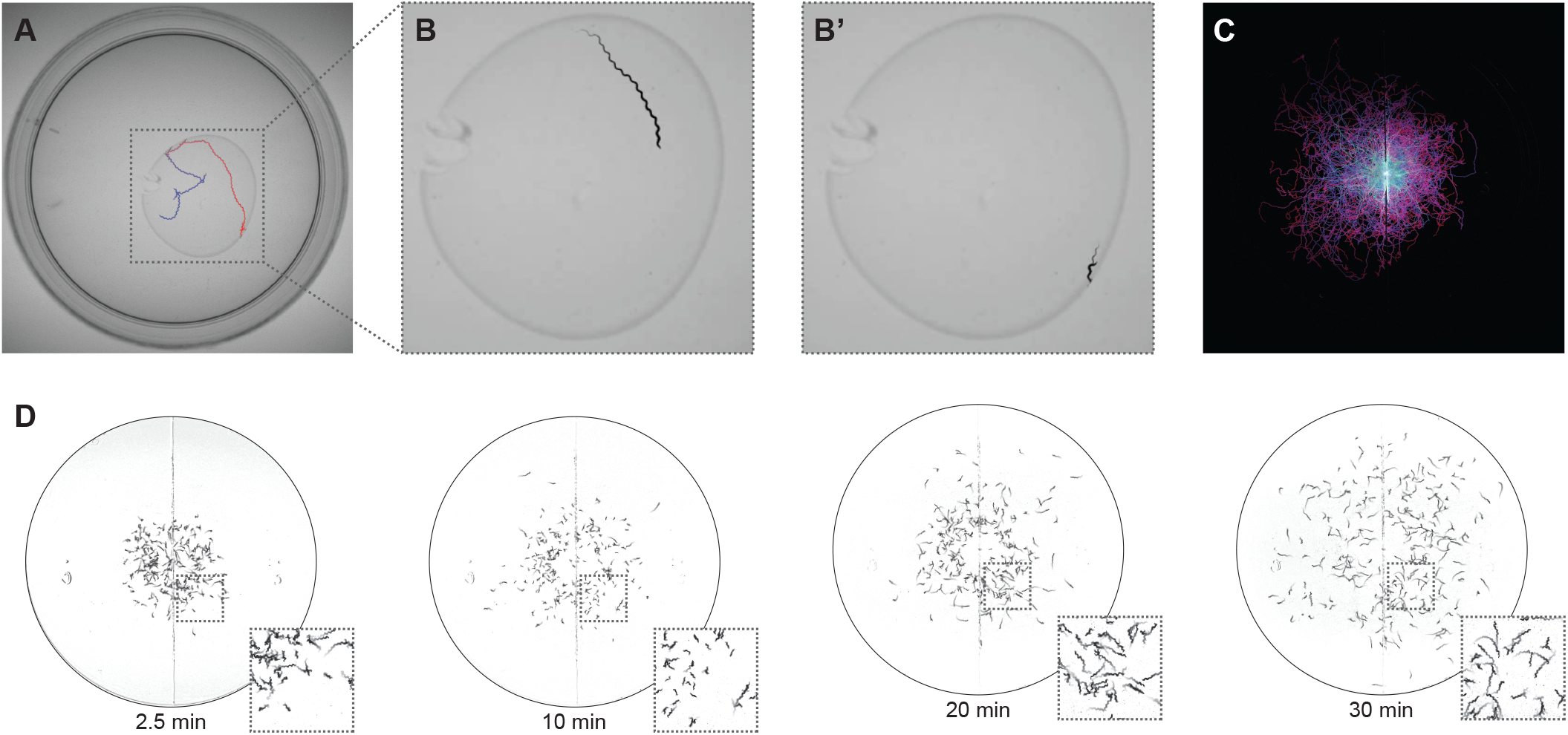
Visualizations of *C. elegans* behavior created using wormtrails. (A) Visualization of the movement of a single worm on a 6 cm petri plate with a bacterial lawn of *E. coli* OP50, over 10 minutes, with earlier times in blue and later times in red. (B-B’) Visualizations showing the locomotion of the worm in panel A over two 60-second windows in which the worm was either (B) roaming or (B′) dwelling in place. (C) Visualization of the first 5 minutes of a chemotaxis assay in a hydrogen peroxide gradient stemming from a point source on the left side of the plate. Trail color transitions from cyan at assay start to magenta at 5 minutes. This visualization received the Best in Microscopy award at the Worm Art Show of the 25th International Worm Meeting. (D) Visualizations of *C. elegans* dispersal on a 9 cm petri plate generated with the default wormtrails settings to show locomotion over the previous 20 seconds at 2.5, 10, 20, and 30 minutes after assay start.

These visualization modes can reveal behavioral dynamics that are difficult to see in raw or average-subtracted videos (compare Videos 1-3 and 4-6). For example, in Figure 1D, worms moved substantially faster 20 minutes into the assay than at 10 minutes, and they tended to follow straighter paths and explore further toward the plate edges. This time-dependent change in speed and exploration was readily apparent in the trails but would be easier to miss in raw or average-subtracted videos.

To support diverse temporal encodings, wormtrails allows user-defined colormaps to be applied to the trails. Several common options are provided, including sequential and divergent colormaps. Sequential color gradients effectively represent locomotory progression from the beginning to the end of the temporal window (Figures 1A,C). In cases with an intervention at a specific timepoint, we favor diverging colormaps to distinguish the locomotory behavior before and after the intervention. The package can generate still images of trails at specific time points and videos of those trails as a function of time (Videos 1-6).

In its current form, wormtrails is well suited for most behavioral assays lasting less than an hour. Future updates will focus on improving performance to support longer and higher-resolution recordings, expanding the library of built-in colormaps, and providing interfaces to common tracking software. A web-based graphical user interface is also under development to make wormtrails accessible to users without a programming background.

## Methods

### Computational techniques

Installation and usage instructions, as well as the full wormtrails source code, are available at https://github.com/ApfeldLab/wormtrails. Most processing steps use the NumPy and OpenCV libraries. The wormtrails processing pipeline consists of two preprocessing steps followed by visualization. Once a video is loaded into a NumPy array, the first preprocessing step is vignetting correction. For each frame, the overall brightness is normalized to a common value and then divided by a Gaussian-blurred average frame to even out low-frequency brightness variation. This mitigates the impact of uneven illumination or poor-quality images, although an ideal imaging setup would minimize the need for vignetting correction. The second preprocessing step is average-frame subtraction, which isolates pixels that change in intensity over the course of the recording. The minimal visualization that can be produced is simply the array of average-subtracted frames written out as a video file. To generate a single-frame projection, the average-subtracted frames are collapsed with a pixel-wise maximum operation, revealing the paths taken by the worms. Each average-subtracted frame is scaled to enhance contrast and offset by a fixed pixel value to remove low-amplitude noise, then multiplied by a color value based on its acquisition time to encode temporal information. For video visualizations, these steps are applied within a moving temporal window across the entire input video.

### Imaging

While most video file formats are compatible with wormtrails, a lossless format is recommended to prevent artifacts. Scans from either bright field or oblique illumination setups will work with wormtrails, because absolute differences are used to compute the average-subtracted frames. To produce continuous trails, frames should be acquired frequently enough that no worm travels more than one body length between successive frames. For most behavioral assays, we recommend an acquisition rate of 1 frame per second. In our experience, trails remain visible even in scans with spatial resolution below one pixel per worm, provided that image noise is sufficiently low. However, resolving detailed trail structure requires a pixel pitch of at most one worm width. For adult *C. elegans* scans, we recommend targeting ∼100 µm per pixel in sample space.

### *C. elegans* culture and strains

*C. elegans* wild-type N2 (Bristol) were cultured according to standard laboratory conditions on nematode growth medium (NGM, 17 g/L agar, 2.5 g/L Bacto Peptone, 3.0 g/L NaCl, 1 mM CaCl_2_, 1 mM MgSO_4_, 25 mM H_2_KPO_4_/HK_2_PO4 pH 6.0, 5 mg/L cholesterol) seeded with *E. coli* OP50 at 20°C.

### Chemotaxis assays

We performed population chemotaxis assays as described (Bargmann & Horvitz, 1991) at 20°C on 9 cm modified chemotaxis agar plates (17 g/L agar, 1 mM CaCl_2_, 1 mM MgSO_4_, 25 mM H_2_KPO_4_/HK_2_PO4 pH 6.0) that we poured 3 days before use and stored at room temperature (20-22°C) (Schiffer et al., 2021). We generated age-synchronous day 1 adults by timed egg laying. We washed these animals three times in M9 buffer and placed them at the center of each chemotaxis plate, and imaged their positions. Images were acquired at 1 frame per second with 1200 x 1200 pixel resolution using a Flir Grasshopper3 USB3 camera positioned vertically over a 9 cm chemotaxis plate, illuminated from below by a Melpo BLFL-LFFA red LED array through a Fisherbrand polystyrene antistatic weighing dish to provide diffuse lighting.

## Figure legends

**Videos 1-3. Single worm locomotion on a 6 cm petri plate with an *E. coli* OP50 bacterial lawn**. (Video 1) Raw video. (Video 2) Average-frame subtracted. (Video 3) Wormtrails visualization showing locomotion over the previous 60 seconds.

**Videos 4-6. Dispersion of a large population of adult worms on a 9 cm petri plate**. (Video 4) Raw video. (Video 5) Average-frame subtracted. (Video 6) Wormtrails visualization showing locomotion over the previous 20 seconds.

## Acknowledgements

We benefitted from discussions with members of Javier Apfeld’s and Erin Cram’s labs. We derived some information from Wormbase, which is supported by the National Human Genome Research Institute at the NIH (grant #U41 HG002223), the UK Medical Research Council, and the UK Biotechnology and Biological Sciences Research Council. Some strains were provided by the CGC, which is funded by NIH Office of Research Infrastructure Programs (P40 OD010440). We acknowledge grant support from NIH grant R21AG086992, a Hevolution Foundation GRO award, and a Longevity Impetus Grant from Norn Group, Hevolution Foundation and Rosenkranz Foundation.

